## Supplemental text and Figure S1-S6. for "Enhanced bacterial chemotaxis in confined microchannels: Optimal performance in lane widths matching circular swimming radius"

### Supplemental materials

**Explanation of difference in drift velocity between LSW and RSW cells.**

We observed obvious chemotaxis of *E. coli* on the surface of lanes, closely related to cells swimming along sidewalls. As shown in Fig. S6A, cells swimming along sidewalls in an attractant gradient can be divided into four states: Cells on the right sidewall swimming up-gradient (RSW-UG), cells on the right sidewall swimming down-gradient (RSW-DG), cells on the left sidewall swimming up-gradient (LSW-UG), and cells on the left sidewall swimming down-gradient (LSW-DG). These cells can escape from sidewalls with rates:

$$k_{RSW-UG} = B^+ k_T$$
$$k_{RSW-DG} = B^- k_T + (1 - B^-) k_R$$
$$k_{LSW-UG} = B^+ k_T + (1 - B^+) k_R$$
$$k_{LSW-DG} = B^- k_T$$

where $B^+$ and $B^-$ represent the probability of tumble when cells swim up-gradient and down-gradient, respectively, and $k_T$ and $k_R$ denote the escape rates during tumble and run, respectively. Cells swimming on the sidewalls experience a force $f_s$ from the bottom surface, causing them to swim to the right. RSW-UG and LSW-DG cells can swim away from the sidewalls via tumble, while RSW-DG and LSW-UG cells can escape via tumble or run. The gradient field direction ensures that $0 \leq B^+ < B^- \leq 1$.

Dwell times for the four states are plotted in Fig. S6B, suggesting that $\tau_{RSW-UG} > \tau_{LSW-DG} > \tau_{RSW-DG} > \tau_{LSW-UG}$. Thus, $k_R > k_T > 0$. This is physically consistent: along sidewalls, running cells escape more easily via circular swimming, whereas tumbling cells are hindered by surface-suppressed reorientation and the high probability of tumbling back toward the sidewalls.

The drift velocity for cells on right and left sidewall can be calculated as:

$$v_d^{RSW} = v_0\left(\frac{1}{k_{RSW-RS}} - \frac{1}{k_{RSW-LS}}\right) = v_0\frac{(B^- - B^+)k_T + (1 - B^-)k_R}{B^+B^-k_T^2 + (1 - B^-)B^+k_Tk_R}$$

$$v_d^{LSW} = v_0\left(\frac{1}{k_{LSW-RS}} - \frac{1}{k_{LSW-LS}}\right) = v_0\frac{(B^- - B^+)k_T - (1 - B^+)k_R}{B^+B^-k_T^2 + (1 - B^+)B^-k_Tk_R}$$

where $v_0$ represents the swimming speed. Given that $0 \le B^+ < B^- \le 1$ and $k_R > k_T > 0$, we can conclude that:

$$v_d^{RSW} > 0$$

$$v_d^{LSW} < 0.$$

**Supplemental movies**

**Movie S1**. An example video of HCB1-pTrc99a-mCherry cells chemotaxis in a 44-µm-wide lane under a linear gradient of L-aspartate. The source channel is on the right side.

**Movie S2.** An example of a large reorientation angle during tumbling in the LSW of a 44-µm-wide lane under a linear gradient of L-aspartate. The source channel is on the right side. The cell of interest is marked with a blue circle.

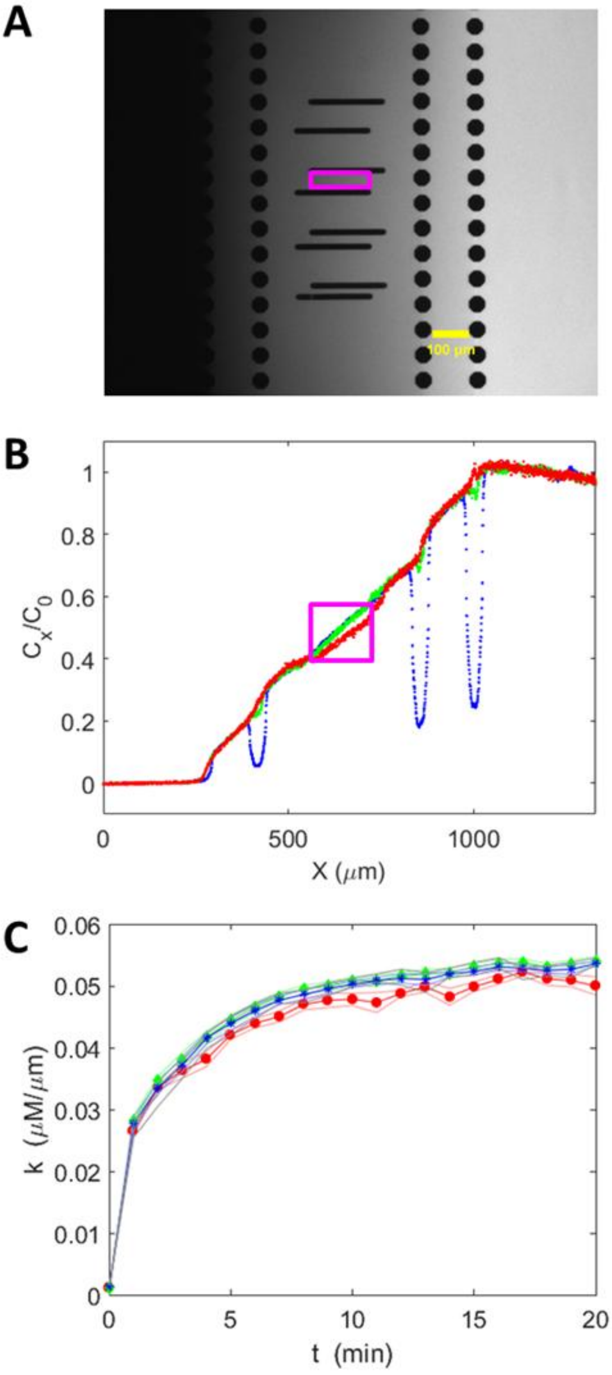


Fig.S1. Gradient calibration of microfluidics with fluorescein **A.** Fluorescence image of the stable gradient field. **B.** Normalized gradient in lanes of different widths. Red, green, and blue lines represent relative concentration values of the substance at different *x* positions for lanes with widths of 15 μm, 25 μm, and 44 μm, respectively. The red rectangle represents the ROI in Fig. S1A. **C.** Change in concentration gradient perceived by bacteria over time in lanes of different widths. Red circles (15 μm), green diamonds (25 μm), and blue asterisks (44 μm) show 11 calibration measurements. Shaded areas represent SEM.

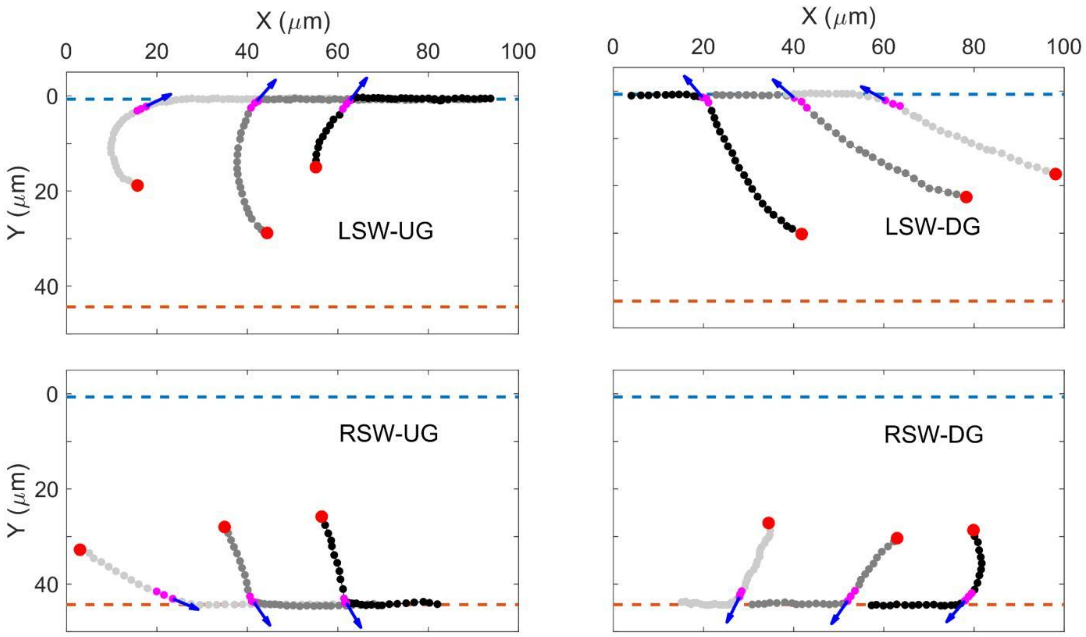


Fig. S2. Typical examples of collision trajectories between bacteria and sidewalls. Data are derived from the 44-μm-wide lane in gradient assays. Typical trajectories (gray dotted lines) illustrate cells colliding with the sidewalls, where different shades of gray represent distinct cells. The left and right sidewalls are indicated by blue and red dashed lines, respectively. Red points denote the starting positions of the trajectories, and blue arrows indicate the direction of the collision.

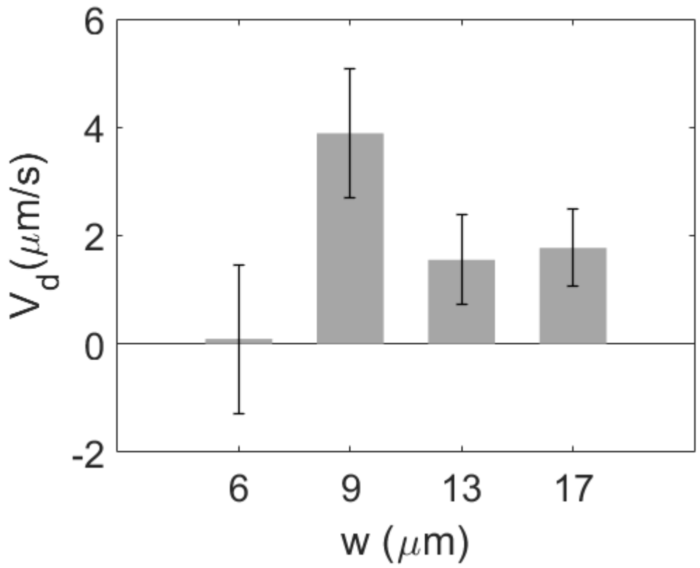

Fig. S3. The relationship between chemotactic drift velocity and channel width measured using a control setup with straight entrances. Channels with widths of 6, 9, 13, and 17 μm contain 217, 238, 434, and 538 trajectories, respectively. Error bars represent the standard deviation of the drift velocity.

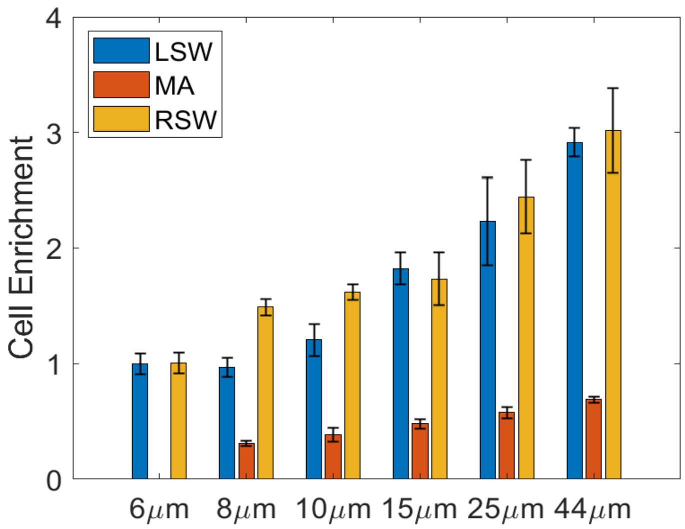


Fig. S4. The relationship between cell enrichment and lane width ($w$). Enrichment values are calculated by as $P\times w/3$ for LSW and RSW, and $P\times w/(w\text{-}6)$ for MA, where $P$ represents the observed cell proportion from Fig. 4B.

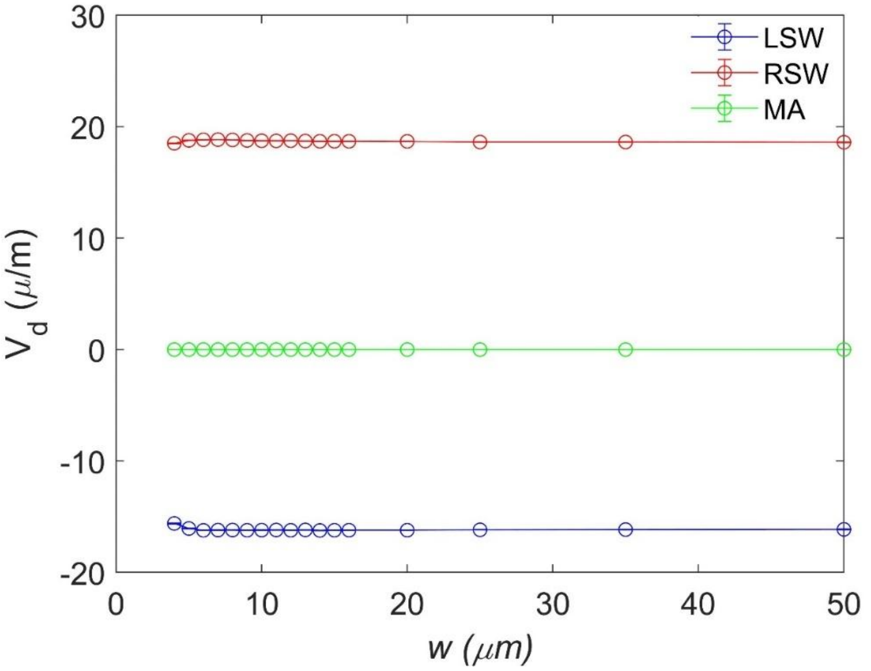


Fig. S5. Mean drift velocities in the LSW, MA, and RSW regions of lanes with different widths from simulations. The simulations assume cells perform a circular swim with a 10 μm radius.

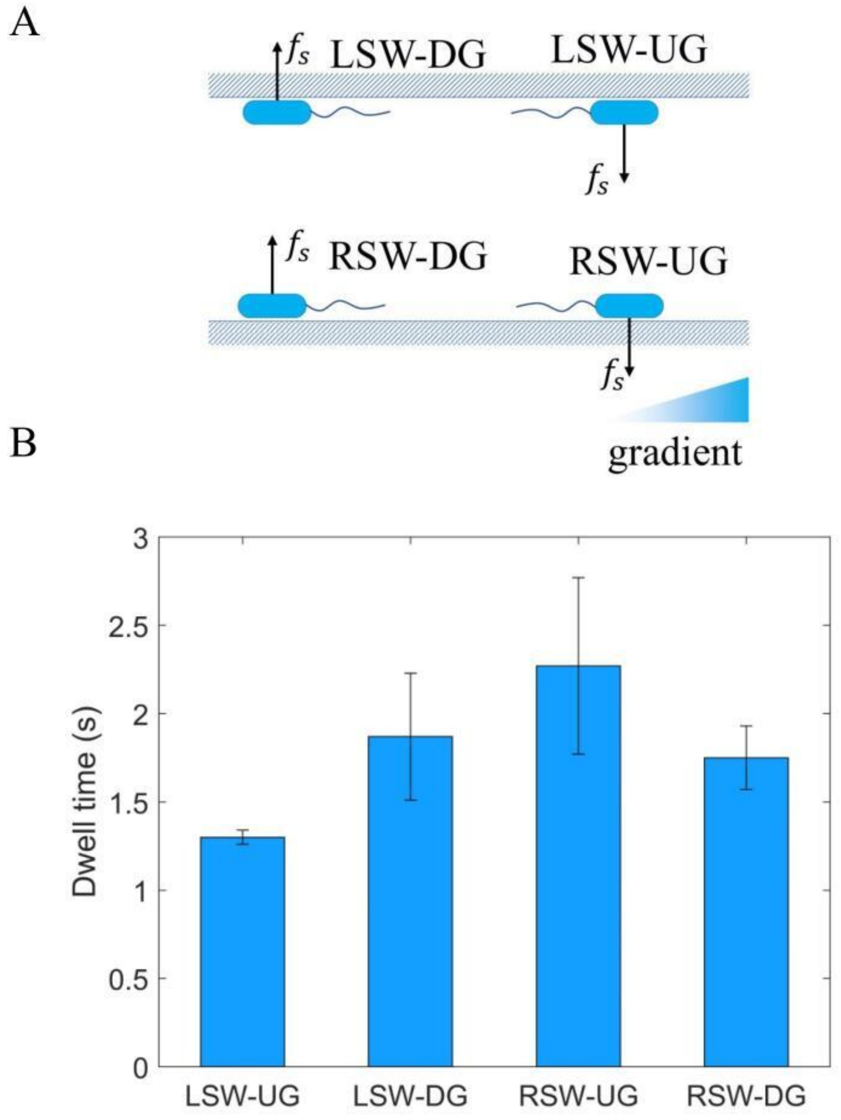


Fig.S6. **A.** Four motion states of cells swimming along sidewalls. Black arrows denote the direction of force $f_s$ generated by the bottom surface on the cell body. **B.** Dwell times of the four motion states from experiments in 44 μm-wide lanes. The values are 1.3±0.04 s, 1.87±0.36 s, 2.27±0.5 s, and 1.75±0.18 s for LSW-UG, LSW-DG, RSW-UG, and RSW-DG respectively. Errors represent SDs.
